# Altered GluN2A levels reduce the emergence of seizure-like events in a rat model of *GRIN2A* haploinsufficiency

**DOI:** 10.1101/2025.06.09.658610

**Authors:** Neela K. Codadu, Giles E. Hardingham, Peter C. Kind, David J. A. Wyllie

**Affiliations:** Centre for Discovery Brain Sciences, Hugh Robson Building, University of Edinburgh, Edinburgh, EH8 9XD, UK; Simons Initiative for the Developing Brain, Hugh Robson Building, University of Edinburgh, Edinburgh, EH8 9XD, UK; UK Dementia Research Institute, University of Edinburgh, Edinburgh, Edinburgh Medical School, Edinburgh, EH16 4SB, UK

**Keywords:** *Grin2a*, GluN2A, NMDA receptor, epilepsy, seizure susceptibilitya

## Abstract

GluN2A-containing NMDA receptors, encoded by the *GRIN2A* gene, are critical components of excitatory synaptic transmission and are essential for proper brain development and function. Dysregulation of NMDA receptor activity is strongly implicated in the pathophysiology of epilepsy. While excessive GluN2A-mediated signalling can lead to hyperexcitability and seizure generation, loss-of-function mutations in *GRIN2A*, which result in reduced GluN2A expression, have also been identified in patients with various forms of epilepsy, including focal epilepsies and epileptic encephalopathies. This paradox highlights the complexity of NMDA receptor contributions to network excitability and suggests that both gain- and loss-of-function mutations can be pathogenic. However, the mechanisms through which reduced GluN2A expression influences seizure susceptibility remains unclear.

In this study, we examined the impact of reduced and absence of GluN2A on seizure-like activity using *in vitro* model system. Hippocampal slices were prepared from wild-type littermate controls (*Grin2a^+/+^*), heterozygous (*Grin2a^+/-^*) and homozygous (*Grin2a^-/-^*) transgenic rats and subjected to pro-convulsant conditions to elicit epileptiform-like events. Specifically, we employed three well established *in vitro* epilepsy models: (i) 4-aminopyridine (4-AP), (ii) zero-magnesium with elevated potassium (0Mg²⁺/5K⁺), and (iii) high-potassium (high-K^+^). Local field potentials were recorded from the CA1 pyramidal layer to quantify interictal and seizure-like activity, by measuring latency to onset, event frequency, and duration. Spectral analyses were also conducted to assess alterations in network dynamics.

Our findings reveal that reduced GluN2A expression alters the susceptibility of hippocampal circuits to seizure-like activity in a model-dependent manner. In the 4-AP model, *Grin2a^-/-^* slices exhibited a significantly lower frequency of interictal events and a delayed onset of seizure-like activity relative to both wild-type and heterozygous slices. In the 0Mg²⁺/5K⁺ model, both *Grin2a^+/-^* and *Grin2a^-/-^* slices displayed reduced seizure susceptibility compared to wild-type. Finally, in the high-K⁺ model, *Grin2a^-/-^* slices showed a reduced incidence of seizure-like events compared to wild-type controls when extracellular potassium concentration was increased to 7.5 mM, although this effect was less apparent at higher concentrations (9.5 mM).

These results suggest that partial or complete loss of GluN2A-containing NMDA receptors reduces the propensity of hippocampal networks to develop epileptiform activity, potentially by altering intrinsic or synaptic excitability. Our findings provide mechanistic insights into how *GRIN2A* loss-of-function variants may contribute to epilepsy and highlight the need to consider developmental and circuit-level compensations when interpreting the impact of GluN2A disruption.

## Introduction

*N*-Methyl-D-aspartate (NMDA) receptors are tetrameric ligand-gated cation channels activated by L-glutamate, the major excitatory neurotransmitter in the mammalian brain. They play critical roles in synaptic transmission, plasticity, cognition, learning, memory, and normal brain development. Majority of NMDA receptors comprise two obligatory GluN1 subunits and any two of the four GluN2 (A-D) subunits.^1–3^ The biophysical properties of NMDA receptors are determined by the GluN2 subunit composition. NMDA receptors containing GluN2A subunits, encoded by *GRIN2A*, either in diheteromeric (GluN1/2A) or triheteromeric (GluN1/2A/2B) configurations, exhibit faster decay kinetics compared to those receptors not containing GluN2A subunits.^1–7^ In addition, GluN2A-NMDA receptor subunits play critical roles in shaping synaptic physiology, and network excitability.^2,8^

Recent genome-wide association studies have identified numerous *GRIN2A* mutations in humans that are strongly associated with a spectrum of epilepsy-aphasia disorders, including Landau-Kleffner syndrome, epilepsy with continuous spikes and waves, and benign epilepsy with centrotemporal spikes.^9–12^ These mutations can result in either gain- or loss-of-function phenotypes.^9,10,13–16^ Loss-of-function *GRIN2A* mutations associated with various forms of epilepsy include missense, nonsense variants, and microdeletions encompassing the *GRIN2A* gene, resulting in null variants that affect receptor properties such as reduced glutamate potencies, altered trafficking, and reduced protein expression.^13,17,18^ The strong genotype–phenotype correlations identified in clinical cohorts suggest a causative role for *GRIN2A*-null variants in epilepsy.

At the cellular level, loss or reduction of GluN2A alters hippocampal CA1 pyramidal neuron synaptic properties, leading to reduced excitatory currents with longer decay, and decreased dendritic arborisation, without significantly affecting intrinsic excitability.^12,19–21^ Given the role of GluN2A in mediating excitatory synaptic transmission, it is generally expected that loss-of-function mutations would reduce excitatory drive and, consequently, lower seizure susceptibility. However, paradoxically, networks with *GRIN2A* loss-of-function mutations can transition into hyperexcitable states that promote seizures. In a mouse model of *GRIN2A* haploinsufficiency, transient epileptiform activity was observed during early development, further highlighting the complexity of GluN2A role in network function.^22^ Although the effects of reduced GluN2A at the cellular-level are well documented, how they translate into altered network-level excitability and epileptiform activity remains unclear. Addressing this gap is crucial for understanding the mechanistic basis of *GRIN2A*-related epilepsies.

In this study, we systematically investigated how the absence or reduction of GluN2A-containing NMDA receptors modulates seizure susceptibility and network-level epileptiform dynamics. To this end, we challenged acute hippocampal slices from *Grin2a* transgenic rats (*Grin2a^+/–^*, and *Grin2a^−/−^*) and wild-type littermate controls (*Grin2a^+/+^*) using three mechanistically distinct *in vitro* models of epilepsy:^23,24^ a) the 4-aminopyridine (4-AP) model, which primarily drives seizures via enhanced interneuronal firing and disinhibition; b) the zero-magnesium/high-potassium (0Mg²⁺/5K⁺) model, which facilitates epileptiform activity through elevated glutamatergic drive; and c) the high-potassium model (high-K^+^), which induces generalised depolarisation across neuronal populations, leading to non-specific network activation. We found that neither heterozygous (*Grin2a^+/-^*) nor homozygous (*Grin2a^-/-^*) *Grin2a* null variants showed increased seizure susceptibility in any of the models tested. Instead, the null variants appeared to demonstrate resistance: *Grin2a^-/-^* displayed increased resistance to the development of seizure-like events (SLEs) in both 4-AP and 0Mg^2+^/5K^+^ models, and *Grin2a^+/-^* showed increased resistance in the 0Mg^2+^/5K^+^ model. In the high-K⁺ model, *Grin2a⁻^/^⁻* displayed a reduced incidence of seizure-like events, although this effect diminished with increasing potassium concentrations. These findings provide novel insights into the complex and context-dependent role of GluN2A-containing NMDA receptors in network excitability and epileptiform activity. Our results suggest that reduced GluN2A does not inherently promote epileptiform activity, and may in some conditions reduce susceptibility to seizures, highlighting the importance of circuit-level adaptations in *GRIN2A*-related seizure phenotypes.

## Materials and methods

### Animals and ethical approval

Animal husbandry and all experiments adhered to the United Kingdom Animals (Scientific Procedures) Act of 1986. Experiments were conducted under the authority of Home Office licence P1351480E and were in accordance with the University of Edinburgh animal welfare committee regulations. The *Grin2a* transgenic rat line, comprising homozygous (*Grin2a^-/-^*), heterozygous (*Grin2a^+/-^*), and wild-type littermates (*Grin2a^+/+^*), was maintained on a Long-Evans Hooded background.^12^ Rats were all housed under a standard 12:12 h light/dark cycle and received food and water *ad libitum*.

### Brain slice preparation

Young-adult rats of age between postnatal days 26 and 38 were sacrificed by non-schedule 1 method. Briefly, rats were anaesthetised using isoflurane, decapitated and their brains rapidly removed and immersed in ice-cold sucrose-based artificial cerebrospinal fluid solution containing (mM): NaCl, 86; NaH_2_PO_4_, 1.2; KCl, 2.5; NaHCO_3_, 25; glucose, 25; sucrose, 75; CaCl_2_, 0.5; MgCl_2_, 7. Hippocampal slices (400 µm thick) were prepared using a vibratome (Leica Biosystems). Slices were then transferred to an interface tissue holding chamber containing artificial cerebrospinal fluid (ACSF) of following composition (mM): NaCl, 124; NaH_2_PO_4_, 1.2; KCl, 2.5; NaHCO_3_, 25; glucose, 20; CaCl_2_, 2; MgCl_2_, 1. Prior to recording, these slices were initially incubated for 30 minutes at 33.5 °C in a water bath (Grant Instruments) followed by 30 minutes at room temperature. All the solutions used for slice preparations and experiments were bubbled continuously to saturate with 95% O_2_ and 5% CO_2_.

### Electrophysiology

Local field potential (LFP) recordings were performed in Hass-type interface recording chamber that was maintained at 33.5-34.5 °C by a chamber heating device. Slices placed in the recording chamber were perfused with ACSF at 3.5-4 mL per minute by a peristaltic pump (Watson-Marlow pumps). Recordings were obtained using borosilicate glass microelectrodes (Harvard Apparatus) pulled using microelectrode puller (P-97, Sutter Instruments) and filled with ACSF. Waveform signals were acquired using an EXT-02B extracellular amplifier (NPI Electronic Instruments), digitised at 10 kHz, amplified (gain:1000) and band-pass filtered (1-5000 Hz) via a BNC-2090 (National Instruments) and acquired using WinEDR (V3.8.6; Strathclyde Electrophysiology Software).

### Induction of epileptiform-like activity

LFPs were recorded from CA1 pyramidal cell layer of the hippocampal slices. Epileptiform-like activity was evoked using following three *in vitro* models of epilepsy: 1) 4-aminopyridine (4-AP) model, ACSF was supplemented with 25, 50, and 100 µM 4-AP; 2) zero-Mg^2+^ + 5 mM K^+^ (0Mg^2+^/5K^+^) model, Mg^2+^ was omitted from the ACSF and supplemented with KCl with the final extracellular K^+^ concentration ([K^+^]_e_) adjusted to 5 mM; 3) high-K^+^ model, ACSF was supplemented with KCl with [K^+^]_e_ adjusted to 5.5, 7.5, and 9.5 mM. LFPs were first recorded for 10-15 minutes in ACSF as baseline followed by 60 minutes in any one of the above pro-epileptic media. In experiments where no activity was observed during the 60 minutes wash with pro-epileptic media, gabazine (GABA_A_ receptor antagonist; 10 µM) was added to the ACSF to test the viability of slices. Only the slices that responded to pro-epileptic media or gabazine were included in the study.

### Data analysis and statistics

*Post hoc* data analysis was performed using Clampfit (V10.5, Molecular Devices), GraphPad, and custom-written scripts in MATLAB R2019b (MathWorks). Power spectral analysis was performed using MATLAB’s built-in function *pwelch* for the frequency bands: delta (1-4 Hz), theta (4-8 Hz), alpha (8-12 Hz), beta (12-25 Hz), gamma (25-48 Hz). Normality of the data was assessed using the Shapiro-Wilk normality test. Datasets with normal distributions were analysed using one-way ANOVA followed by Tukey’s multiple comparisons test. Non-normally distributed data were analysed using the Kruskal-Wallis test followed by Dunn’s multiple comparisons test. The rates of interictal events (IIE) at different concentrations of 4-AP within a genotype showing normal distributions were compared using paired t-test or else by Wilcoxon matched-pairs signed rank test. Repeated measures one-way ANOVA was used to test the rates of interictal event measures taken at different time points within a genotype. Data are presented as mean ± SD and ‘n’ values represent the number of rats per group, unless mentioned otherwise.

### Terminology

In this study, we use the terms (i) seizure-like events (SLEs) to refer to electrophysiological signals that resemble tonic-clonic seizure events, (ii) interictal evets (IIEs) to refer to transient rhythmic discharges, and (iii) ‘epileptiform-like events’ as an umbrella term to refer to both seizure-like events and interictal events.

## Results

### *Grin2a^-/-^* show reduced network excitability and increased latency to SLEs in the 4-AP model

4-Aminopyridine (4-AP) is known to cause hyperexcitability and induce seizure activity both *in vitro* and *in vivo*. A hallmark of 4-AP-induced epileptiform activity is the presence of transient recurrent discharges, commonly referred to as interictal events (IIEs).^25–27^ We first investigated the effect of reduced GluN2A levels (*Grin2a^+/-^* and *Grin2a^-/-^*) on the development of epileptiform-like (SLEs and IIEs) activity using low concentrations of 4-AP (25 µM and 50 µM). At these concentrations, IIEs, but not SLEs, were developed in slices prepared from *Grin2a^+/+^, Grin2a^+/-^* and *Grin2a^-/-^* rats (Fig. 1 Ai-Ci). At 25 µM 4-AP, IIEs occurred at a lower rate that was consistent across all genotypes. Raising the 4-AP concentration to 50 µM significantly increased the rate of IIEs for each genotype (Fig. 1 Aii-Cii). However, comparing the genotypes revealed that IIE rates in *Grin2a^+/-^* were similar to wild-type at both 4-AP concentrations, whereas *Grin2a^-/-^* showed significantly lower rates at 50 µM 4-AP (Fig. 1D).

**Figure. 1.**
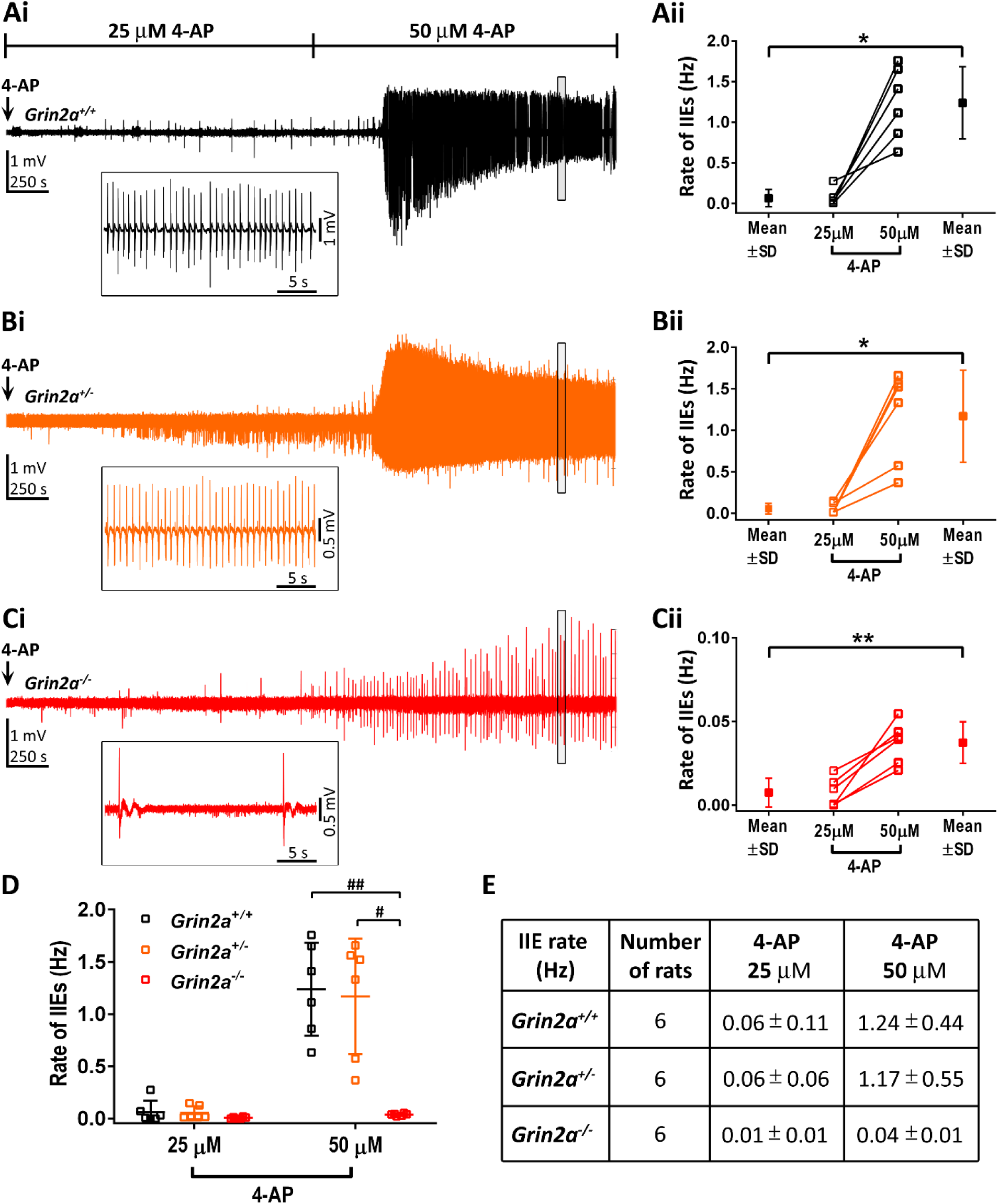
*Grin2a^-/-^* show reduced network excitability with fewer 4-AP-induced interictal events. **(Ai-Ci)** Representative local field potential (LFP) recordings of interictal events induced by bathing (Ai) *Grin2a^+/+^,* (Bi) *Grin2a^+/-^* and (Ci) *Grin2a^-/-^* slices in 25 and 50 μM 4-AP. Insets: Box highlights of interictal events on faster timescales. **(Aii-Cii)** Quantification of the rate of interictal events developed at 25 and 50 μM 4-AP within genotypes: (Aii) *Grin2a^+/+^,* Wilcoxon matched-pairs signed rank test, p = 0.03; (Bii) *Grin2a^+/-^*, Wilcoxon matched-pairs signed rank test, p = 0.03; (Cii) *Grin2a^-/-^*, paired t-test, p = 0.002. **(D)** Comparison of rate of IIEs between genotypes revealed lower network excitability of *Grin2a^-/-^* in 50 µM 4-AP ACSF (25 µM 4-AP: Kruskal-Wallis statistic = 3.23, p = 0.21; 50 µM 4-AP: Kruskal-Wallis statistic = 11.42, p < 0.001; Dunn’s test: *Grin2a^+/+^* vs *Grin2a^+/-^*, p > 0.99; *Grin2a^+/+^* vs *Grin2a^-/-^*, p = 0.007; *Grin2a^+/-^* vs *Grin2a^-/-^*, p = 0.01). **(E)** Mean±SD values for rate of interictal events measured in 25 and 50 µM 4-AP.

Supplementing ACSF with 4-AP (100 µM) produced robust SLEs and IIEs in all the genotypes (Fig. 2A; Supplementary Table 1A). Recurrent IIEs developed in all the genotypes, and their rates were consistent between genotypes at all three time points tested (Fig. 2E; Supplementary Table 1A). The latency of SLEs in *Grin2a^+/-^* was similar to *Grin2a^+/+^*, whereas it was longer in *Grin2a^-/-^* (Fig. 2B). However, in both *Grin2a^+/-^* and *Grin2a^-/-^* slices once the SLEs were initiated, the pattern of activity progression, duration, and number of SLEs developed were similar to *Grin2a^+/+^* (Fig. 2C, D). Power spectral analysis of SLEs recorded in *Grin2a^+/-^* and *Grin2a^-/-^* revealed no significant differences compared to *Grin2a^+/+^*, either in the power contained in the lower (delta, theta, and alpha; Fig. 3D, Ei) or higher frequency bands (beta, gamma; Fig. 3D, Eii; Supplementary Table 1B). In summary, *Grin2a^-/-^* slices showed resistance to developing 4-AP-induced SLEs, whereas SLEs in *Grin2a^+/-^* slices were similar to those in *Grin2a^+/+^* for the parameters examined. Moreover, at low concentrations of 4-AP (50 µM), *Grin2a^-/-^* slices exhibited a lower rate of IIEs indicating reduced network excitability.

**Figure. 2.**
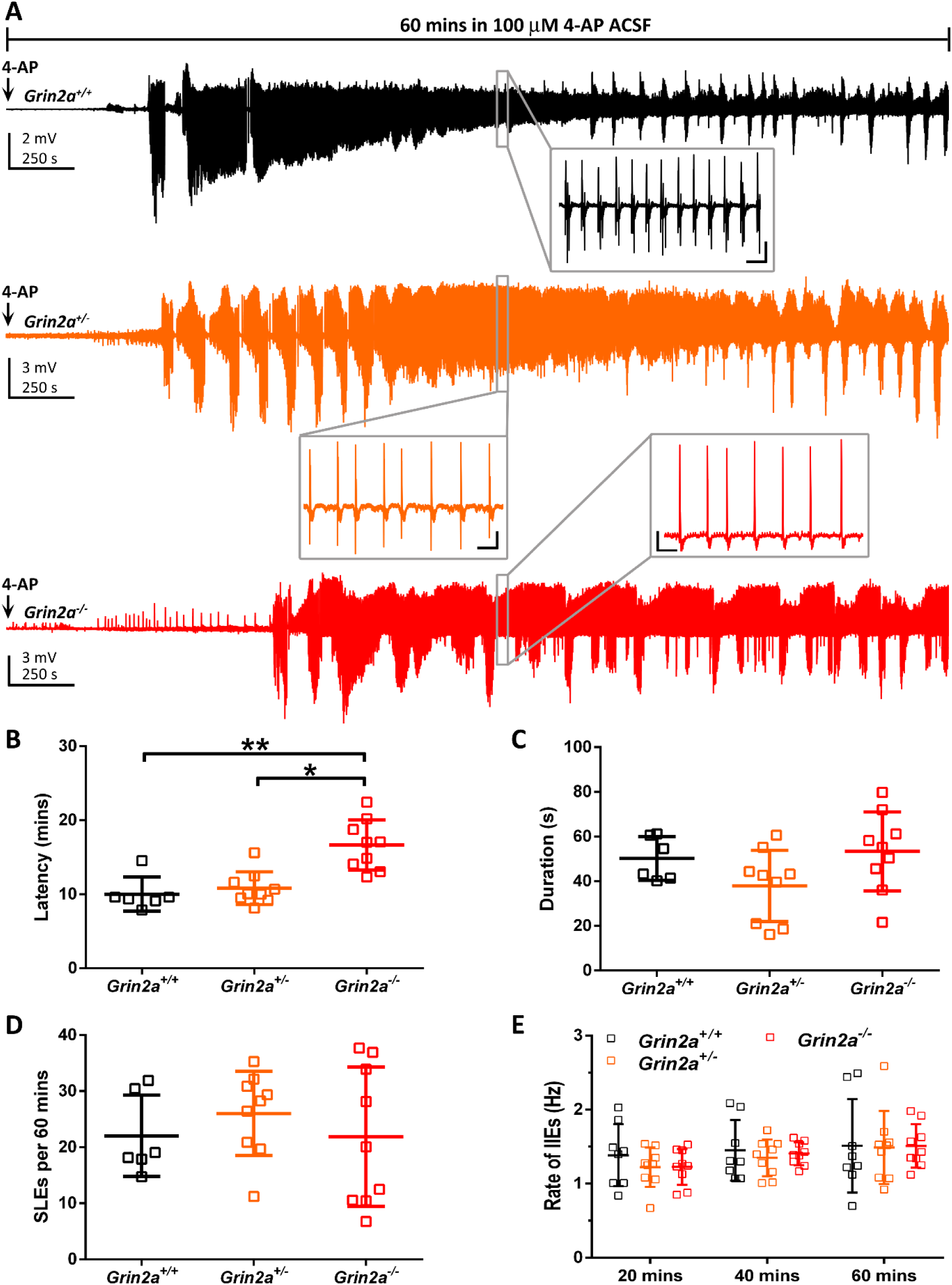
*Grin2a^-/-^* are more resistant for the development of seizure-like events in 100 μM 4-AP, and *Grin2a^+/-^* were similar to wild-types (*Grin2a^+/+^*). (A) Representative local field potential (LFP) recordings from *Grin2a^+/+^, Grin2a^+/-^* and *Grin2a^-/-^* slices bathed in ACSF supplemented with 100 μM 4-AP for 1 hour. Insets: Box highlights of interictal events on faster timescales. Scalebars (genotype, x-axis and y-axis;): *Grin2a^+/+^*, 1 sec and 0.4 mV; *Grin2a^+/-^*, 1 sec and 0.5 mV; *Grin2a^-/-^*, 1 sec and 0.5 mV. **(B)** Quantification of latency to the development of first SLE: *Grin2a^+/+^*, 10.02 ± 2.30 mins (n=6)*; Grin2a^+/-^*, 10.82 ± 2.21 mins (n=9); *Grin2a^-/^*^-^, 16.65 ± 3.39 mins (n=9); Kruskal-Wallis statistic = 3.17, p = 0.001; Dunn’s test: *Grin2a^+/+^* vs *Grin2a^+/-^*, p > 0.99; *Grin2a^+/+^* vs *Grin2a^-/-^*, p = 0.003; *Grin2a^+/-^* vs *Grin2a^-/-^*, p = 0.01. **(C)** Quantification of duration of SLEs: *Grin2a^+/+^*, 50.20 ± 9.75 s (n=6)*; Grin2a^+/-^*, 37.93 ± 15.96 s (n=9); *Grin2a^-/^*^-^, 53.3 ± 17.73 s (n=9); One-way ANOVA, F_2,21_ = 2.43, p = 0.11. **(D)** Quantification of number of SLEs in 60 mins: *Grin2a^+/+^*, 22.03 ± 7.25 (n=6)*; Grin2a^+/-^*, 26.02 ± 7.49 (n=9); *Grin2a^-/^*^-^, 21.88 ± 12.45 (n=9); One-way ANOVA, *F*_2,21_ = 0.50, p = 0.61. **(E)** Quantification of rate of interictal events at three different timepoints after washing in 100 μM 4-AP ACSF; 20 mins: *Grin2a^+/+^*, 1.39 ± 0.42 Hz (n=8)*; Grin2a^+/-^*, 1.22 ± 0.27 Hz (n=9); *Grin2a^-/^*^-^, 1.23 ± 0.24 Hz (n=9); One-way ANOVA, *F*_2,23_ = 0.71, p = 0.50; 40 mins: *Grin2a^+/+^*, 1.45 ± 0.41 Hz (n=8)*; Grin2a^+/-^*, 1.35 ± 0.25 Hz (n=9); *Grin2a^-/^*^-^, 1.40 ± 0.15 Hz (n=9); One-way ANOVA, *F*_2,23_ = 0.28, p = 0.75; 60 mins: *Grin2a^+/+^*, 1.51 ± 0.63 Hz (n=8)*; Grin2a^+/-^*, 1.49 ± 0.49 Hz (n=9); *Grin2a^-/^*^-^, 1.51 ± 0.29 Hz (n=9); One-way ANOVA, *F*_2,23_ = 0.005, p = 0.99. Repeated measures one-way ANOVA determined that rate of IIEs did not significantly change with time (20, 40, and 60 minutes after challenging with 4-AP): *Grin2a^+/^*^+^: *F*_1.14, 7.98_ = 0.57, p = 0.491, n=8; *Grin2a^+/^*^-^: *F*_1.31, 10.44_ = 2.27, p = 0.159, n=9; *Grin2a^-/^*^-^: *F*_1.21, 9.73_ = 4.68, p = 0.051, n=9. For multiple comparisons, see Supplementary Table 1A.

**Figure. 3.**
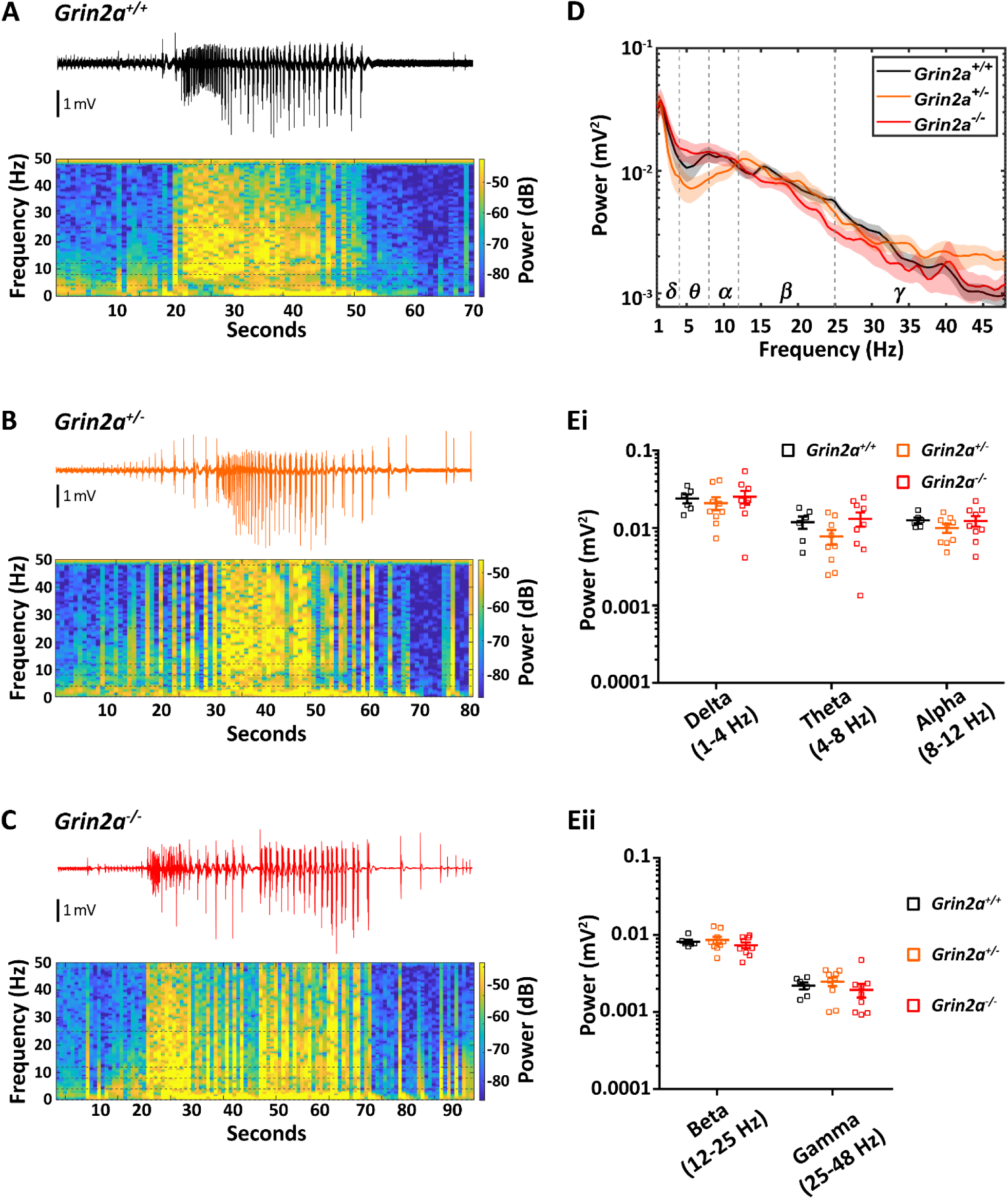
Spectral analysis of 4AP-induced SLEs (100 μM) reveals no differences between *Grin2a^+/-^*, *Grin2a^-/-^* and wild-type (*Grin2a^+/+^*) littermates. (A-C) Representative local field potential (LFP) recordings and associated spectrograms of SLEs from (A) *Grin2a^+/+^,* (B) *Grin2a^+/-^* and (C) *Grin2a^-/-^*. **(D)** Mean power spectral traces for each genotype (solid line: mean; shading: sem). **(Ei, Eii)** Quantification of power in different frequency bands. Delta: One-way ANOVA, F_2,21_ = 0.31, p = 0.74; Theta: One-way ANOVA, F_2,21_ = 1.63, p = 0.22; Alpha: One-way ANOVA, F_2,21_ = 0.86, p = 0.44; Beta: One-way ANOVA, F_2,21_ = 0.83, p = 0.45; Gamma: One-way ANOVA, F_2,21_ = 0.71, p = 0.51. For mean values and multiple comparisons, see Supplementary Table 1B.

### *Grin2a^+/-^* and *Grin2a^-/-^* show reduced seizure susceptibility in the 0Mg^2+^/5K^+^ model

Next, we employed the 0Mg^2+^/2.5K^+^ model to assess seizure susceptibility in the expectation that omission of Mg^2+^ from the ACSF would remove the Mg^2+^ voltage-dependent block of NMDA receptors and increase spontaneous network activity leading to the generation of SLEs.^23,28^ However, no SLEs or any type of epileptiform activity were observed with the 0Mg^2+^/2.5K^+^ model in all genotypes examined until the further addition of gabazine (Supplementary Fig. 1) which had the additional effect of confirming the viability of the slices to generate such events.

To increase the probability of generating SLEs using 0Mg^2+^ model, we increased [K^+^]_e_ from 2.5 to 5 mM.^29^ This concentration of [K^+^]_e_ itself does not elicit epileptiform-like discharges (see Supplementary Fig. 2). The 0Mg^2+^/5K^+^ model induced robust SLEs across all genotypes (Fig. 4A). However, SLEs occurred with a significantly longer latency in both *Grin2a^+/-^* and *Grin2a^-/-^* relative to *Grin2a^+/+^* (Fig. 4B; Supplementary Table 2A). The duration of SLEs in *Grin2a^+/-^* and *Grin2a^-/-^* displayed a trend towards lower values, though not statistically significant (Fig. 4C; Supplementary Table 2A). Additionally, the number of SLEs observed was significantly lower, and the inter-event intervals were significantly higher in *Grin2a^-/-^* (Fig. 4D, E). *Grin2a^+/-^* showed a non-significant trend towards a decrease in number of SLEs and an increase in inter-event intervals (Fig. 4D, E; Supplementary Table 2A).

**Figure. 4.**
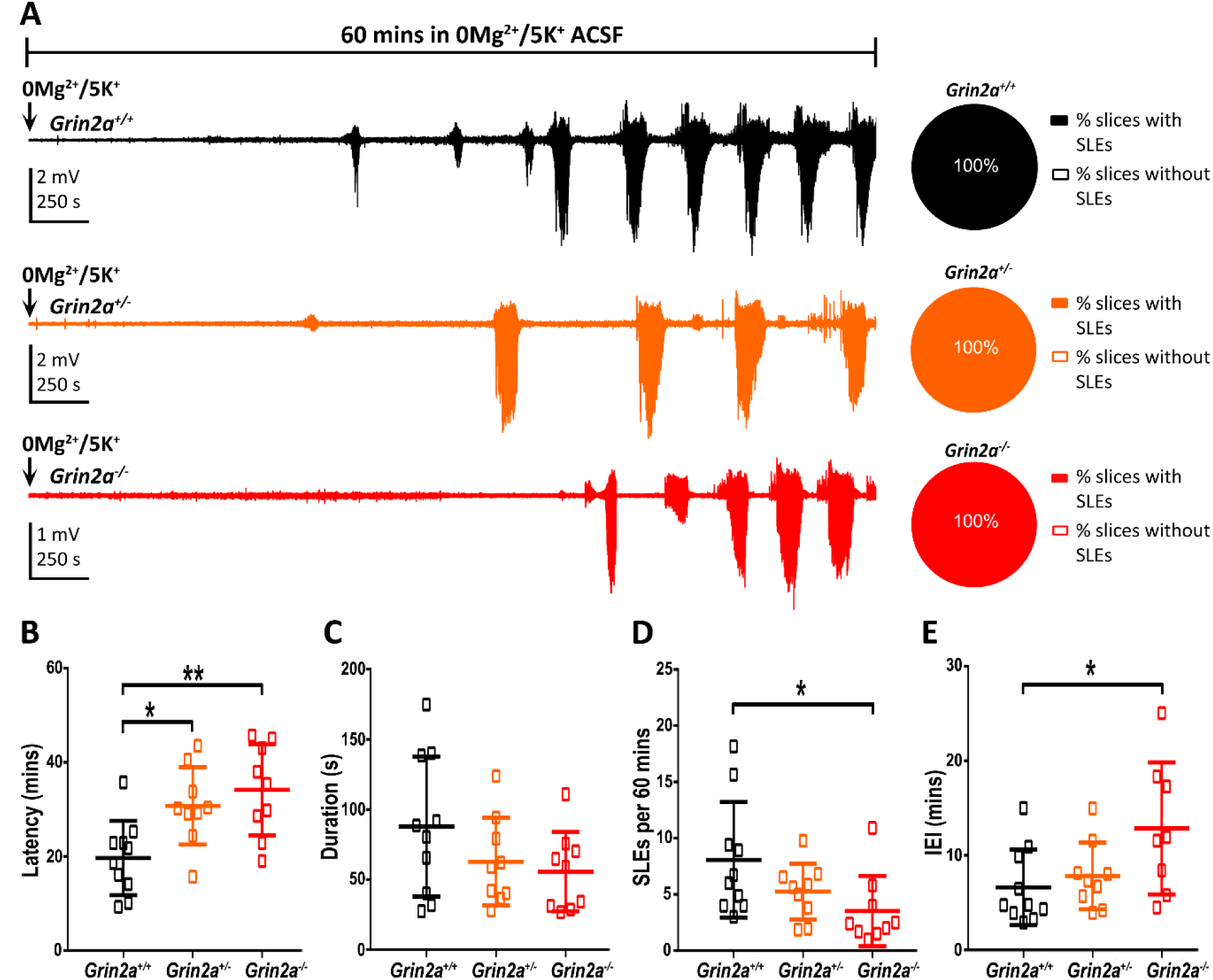
*Grin2a^+/-^* and *Grin2a^-/-^* are relatively resistant to the development of SLEs in 0Mg^2+^/5K^+^ model. **(A)** Left: Representative LFP recordings from *Grin2a^+/+^, Grin2a^+/-^* and *Grin2a^-/-^* slices bathed in zero-magnesium + 5 mM potassium (0Mg^2+^/5K^+^) ACSF for 60 minutes. Right: Pie graph representing percentage of slices with and without SLEs. Number of slices with SLEs/total number of slices (n rats): *Grin2a^+/+^*, 12/12 (10); *Grin2a^+/-^*, 9/9 (9 rats); *Grin2a^-/-^*, 13/13 (9 rats). **(B)** Quantification of latency to the development of first SLE: *Grin2a^+/+^*, 19.69 ± 7.90 mins (n=10)*; Grin2a^+/-^*, 30.77 ± 8.22 mins (n=9); *Grin2a^-/^*^-^, 34.18 ± 9.71 mins (n=9); One-way ANOVA, F_2,25_ = 7.42, p = 0.003; Tukey’s test: *Grin2a^+/+^* vs *Grin2a^+/-^*, p = 0.026; *Grin2a^+/+^* vs *grin2a^-/-^*, p = 0.003; *Grin2a^+/-^* vs *Grin2a^-/-^*, p = 0.683. **(C)** Quantification of duration of SLEs: *Grin2a^+/+^*, 88.96 ± 50.03 s (n=10)*; Grin2a^+/-^*, 62.82 ± 31.30 s (n=9); *Grin2a^-/^*^-^, 55.72 ± 28.26 s (n=9); One-way ANOVA *F*_2,25_ = 1.88, p = 0.17. **(D)** Quantification of number of SLEs in 60 mins: *Grin2a^+/+^*, 8.07 ± 5.14 (n=10)*; Grin2a^+/-^*, 5.25 ± 5.14 (n=9); *Grin2a^-/^*^-^, 3.53 ± 3.12 (n=9); Kruskal-Wallis statistic = 6.76, p = 0.03; Dunn’s test: *Grin2a^+/+^* vs *Grin2a^+/-^*, p = 0.905; *Grin2a^+/+^* vs *Grin2a^-/-^*, p = 0.028; *Grin2a^+/-^* vs *Grin2a^-/-^*, p = 0.386. **(E)** Quantification of inter-event intervals (IEI): *Grin2a^+/+^*, 6.62 ± 3.99 mins (n=10)*; Grin2a^+/-^*, 7.84 ± 3.52 mins (n=9); *Grin2a^-/^*^-^, 12.86 ± 6.97 mins (n=9); Kruskal-Wallis statistic = 6.19, p = 0.62; Dunn’s test: *Grin2a^+/+^* vs *Grin2a^+/-^*, p > 0.99; *Grin2a^+/+^* vs *Grin2a^-/-^*, p = 0.041; *Grin2a^+/-^* vs *Grin2a^-/-^*, p = 0.329. For multiple comparisons, see Supplementary Table 2A.

Power spectral analysis of SLEs revealed a distinctive profile for these events in *Grin2a^-/-^* characterised by reduced power in the lower frequency bands and increased power in the gamma band (Fig. 5A-C, D). A significant reduction in power was observed across delta, theta, and alpha bands (Fig. 5Ei; Supplementary Table 2B). In the higher frequency range, beta band power was similar across genotypes, whereas gamma band power was significantly increased in *Grin2a^-/-^* (Fig. 5Eii; Supplementary Table 2B). Amplified gamma band oscillations have recently been reported in *Grin2a*-deficient networks^30^. Consistent with these findings, our data show increased gamma band power in *Grin2a^-/-^*, which may reflect altered inhibitory network dynamics and a dysregulated inhibitory tone that facilitates gamma oscillations. In summary, *Grin2a^-/-^* was more resistant to developing SLEs in 0Mg^2+^/5K^+^ ACSF, and the SLEs that did occur exhibited distinct spectral features, suggesting altered network synchronisation and excitability.

**Figure. 5.**
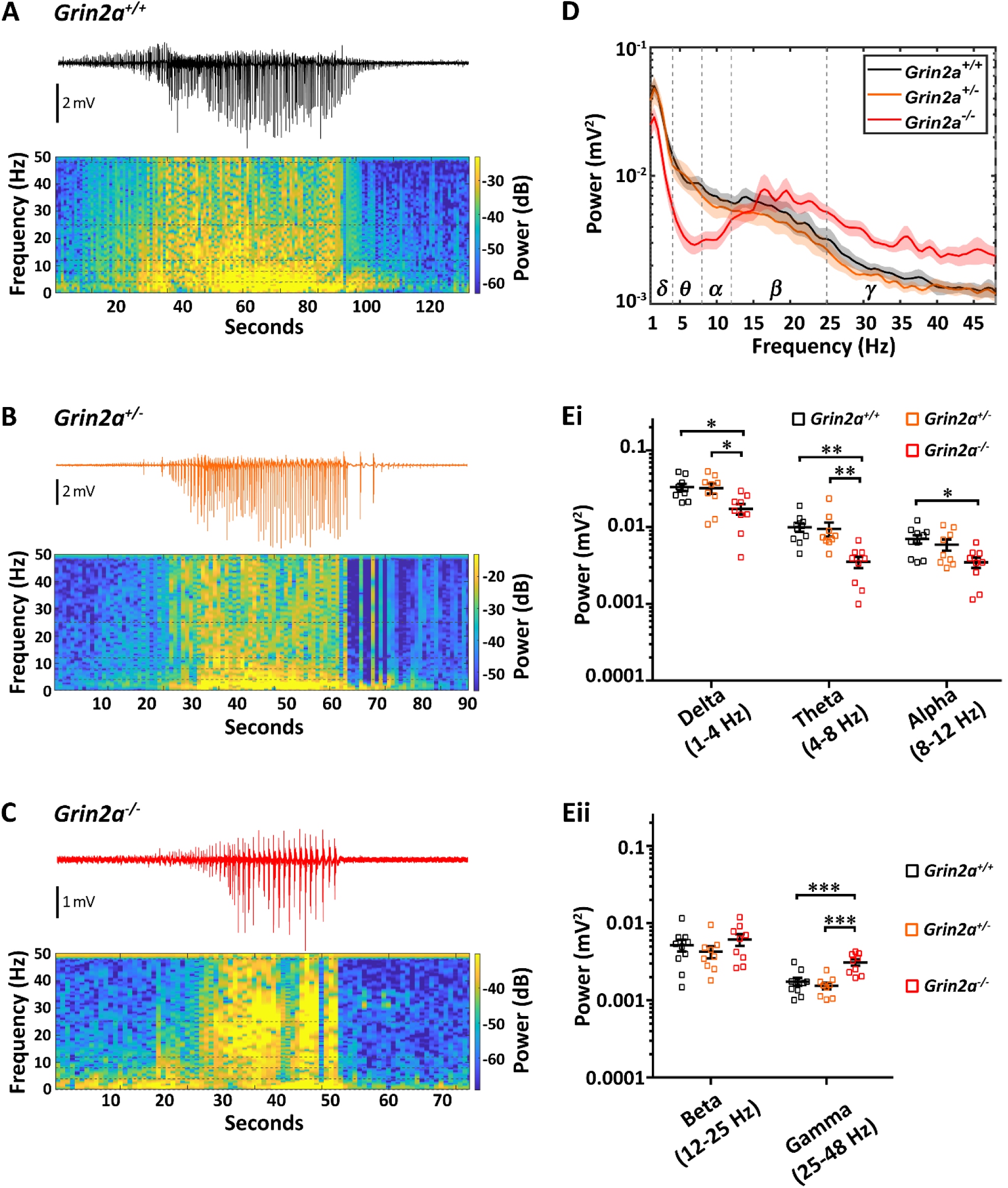
Spectral analysis of SLEs induced by 0Mg^2+^/5K^+^ ACSF reveals distinct spectral features in *Grin2a^-/-^* compared to *Grin2a^+/-^* and wild-type (*Grin2a^+/+^*) littermates. (A-C) Representative LFP recordings and associated spectrograms of 0Mg^2+^/5K^+^ ACSF-induced SLEs in (A) *Grin2a^+/+^,* (B) *Grin2a^+/-^* and (C) *Grin2a^-/-^*. **(D)** Mean power spectral traces for each genotype (solid line: mean; shading: sem). **(Ei, Eii)** Quantification of power in different frequency bands. Delta: One-way ANOVA *F*_2,25_ = 5.65, p = 0.009; Theta: Kruskal-Wallis statistic = 14.46, p < 0.001; Alpha: One-way ANOVA *F*_2,25_ = 4.53, p = 0.02; Beta: One-way ANOVA *F*_2,25_ = 1.04, p = 0.36; Gamma: One-way ANOVA *F*_2,25_ = 13.75, p < 0.0001. For mean values and multiple comparisons, see Supplementary Table 2B.

### *Grin2a^+/-^* and *Grin2a^-/-^* develop seizure-like events similar to littermate controls (*Grin2a^+/+^*) in the high-K^+^ model

Increasing extracellular potassium ion concentrations has been shown to cause the development of SLEs that are sensitive to NMDA receptor antagonists in hippocampal slices.^24^ We first investigated whether there was a difference in the extracellular K^+^ concentration threshold for the development of SLEs in *Grin2a^+/-^* and *Grin2a^-/-^*. Raising the extracellular K^+^ concentration ([K^+^]_e_) to 5.5 mM did not evoke epileptiform-like discharges in any of the genotypes examined (Supplementary Fig. 2). Whereas, 7.5 mM [K^+^]_e_ ACSF evoked SLEs, but in only a proportion of slices from all genotypes. Furthermore, we found that *Grin2a^-/-^* was less likely to develop SLEs (Fig. 6B). When challenged with 9.5 mM [K^+^]_e_ ACSF, all genotypes displayed robust SLEs (Fig. 7A). The latencies, durations, number of SLEs, and inter-event intervals (IEI) were all consistent across *Grin2a^+/+^, Grin2a^+/-^* and *Grin2a^-/-^*genotypes (Fig. 7B-E; Supplementary Table 3A). Power spectral analysis of SLEs indicated that the spectral profiles of *Grin2a^+/-^* and *Grin2a^-/-^* SLEs were similar to those of wild-type controls across all the frequency bands examined (Fig. 8; Supplementary Table 3B). In summary, we observed that at a high extracellular potassium ion concentration (9.5 mM), *Grin2a* null variants showed seizure susceptibilities comparable to wild-types. Whereas, at 7.5 mM [K^+^]_e_, *Grin2a^-/-^* slices were less likely to develop SLEs, and at lower concentrations (5.5 mM [K^+^]_e_), no epileptiform-like activity was observed in either *Grin2a* null variants or wild-types.

**Figure. 6.**
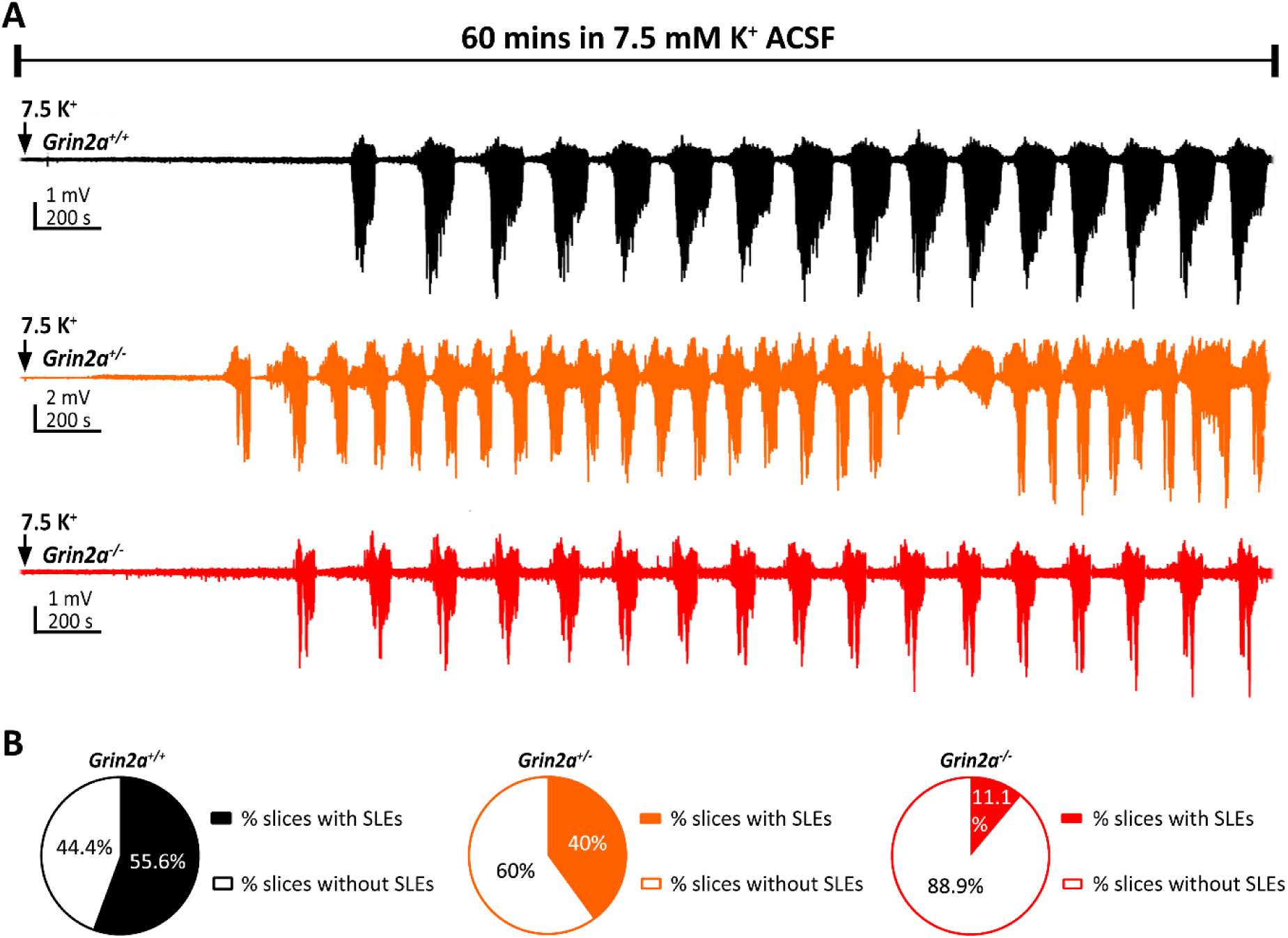
*Grin2a^+/-^* and *Grin2a^-/-^* are less likely to develop SLE in 7.5 mM extracellular potassium levels. **(A)** Representative LFP recordings from *Grin2a^+/+^, Grin2a^+/-^* and *Grin2a^-/-^* slices bathed in ACSF containing 7.5 mM potassium ion concentration for 60 minutes. **(B)** Pie graph representing percentage of slices with and without SLEs. Number of slices with SLEs/total number of slices (n rats): *Grin2a^+/+^*, 5/9 slices (6 rats); *Grin2a^+/-^*, 4/10 slices (6 rats); *Grin2a^-/-^*, 1/9 slices (6 rats); Chi-square test: *Grin2a^+/+^* vs *Grin2a^+/-^*, p = 0.49; *Grin2a^+/+^* vs *Grin2a^-/-^*, p = 0.04; *Grin2a^+/-^* vs *Grin2a^-/-^*, p = 0.15.

**Figure. 7.**
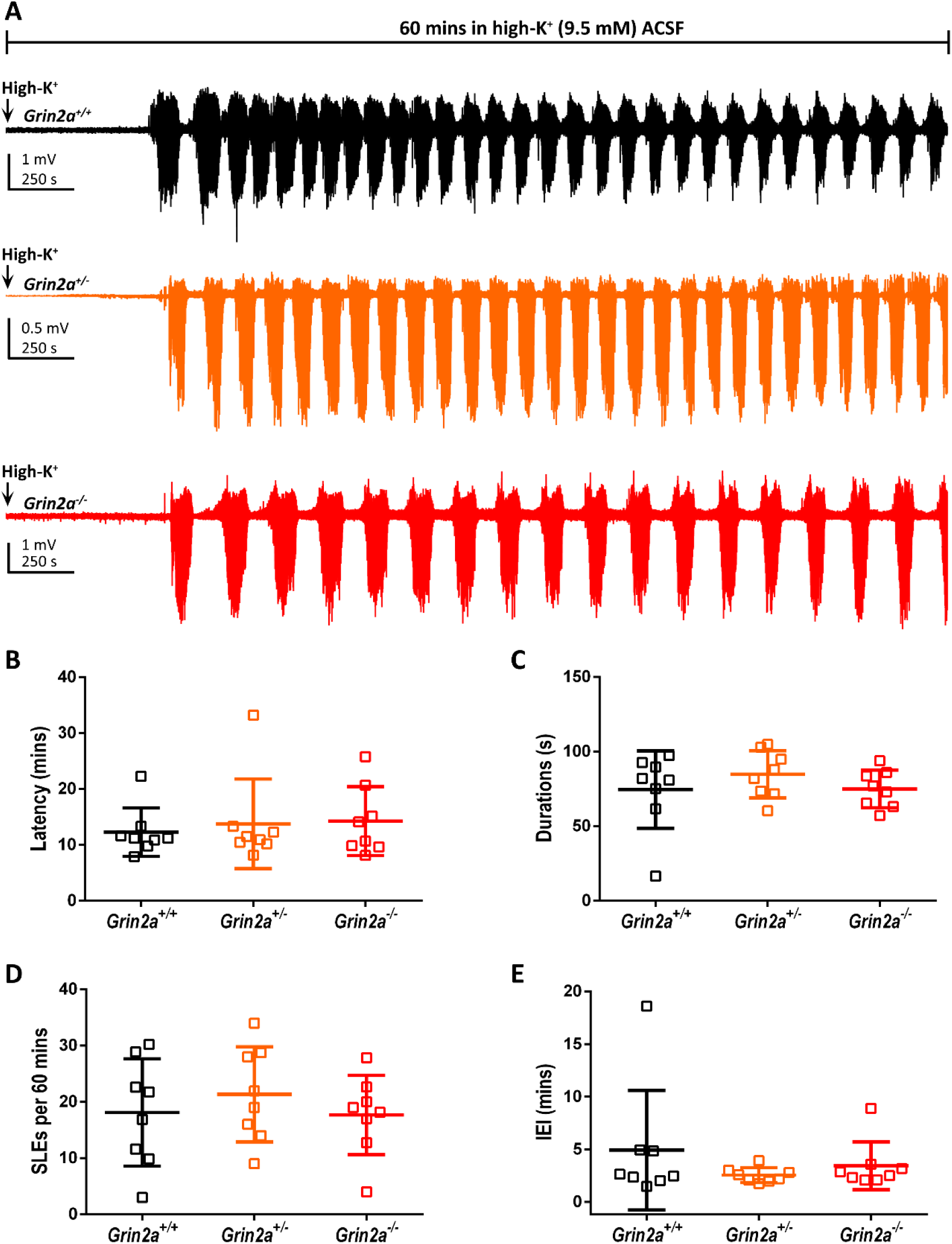
*Grin2a* null variants (*Grin2a^+/-^* and *Grin2a^-/-^*) developed seizure-like events similar to littermate wild-types (*Grin2a^+/+^*) at 9.5 mM extracellular potassium levels. (A) Representative LFP recordings from *Grin2a^+/+^, Grin2a^+/-^* and *Grin2a^-/-^* slices bathed in ACSF containing 9.5 mM potassium ion levels ([K^+^]_e_) for 1 hour. **(B)** Quantification of latency to the development of first SLE: *Grin2a^+/+^*, 12.28 ± 4.33 mins (n=8)*; Grin2a^+/-^*, 13.76 ± 8.01 mins (n=8); *Grin2a^-/^*^-^, 14.25 ± 6.16 mins (n=8); Kruskal-Wallis statistic = 0.11, p = 0.95. **(C)** Quantification of duration of SLEs: *Grin2a^+/+^*, 74.51 ± 25.91 s (n=8)*; Grin2a^+/-^*, 84.77 ± 15.82 s (n=8); *Grin2a^-/^*^-^, 74.91 ± 12.59 s (n=8); Kruskal-Wallis statistic = 1.45, p = 0.49. **(D)** Quantification of number of SLEs in 60 mins: *Grin2a^+/+^*, 18.11 ± 9.53 (n=8)*; Grin2a^+/-^*, 21.34 ± 8.45 (n=8); *Grin2a^-/^*^-^, 17.68 ± 7.04 (n=8); One-way ANOVA, F_2,21_ = 1.45, p = 0.64. **(E)** Quantification of inter-event intervals: *Grin2a^+/+^*, 4.91 ± 5.68 mins (n=8)*; Grin2a^+/-^*, 2.53 ± 0.71 mins (n=8); *Grin2a^-/^*^-^, 3.43 ± 2.26 mins (n=8); Kruskal-Wallis statistic = 0.97, p = 0.62. For multiple comparisons, see Supplementary Table 3A.

**Figure. 8.**
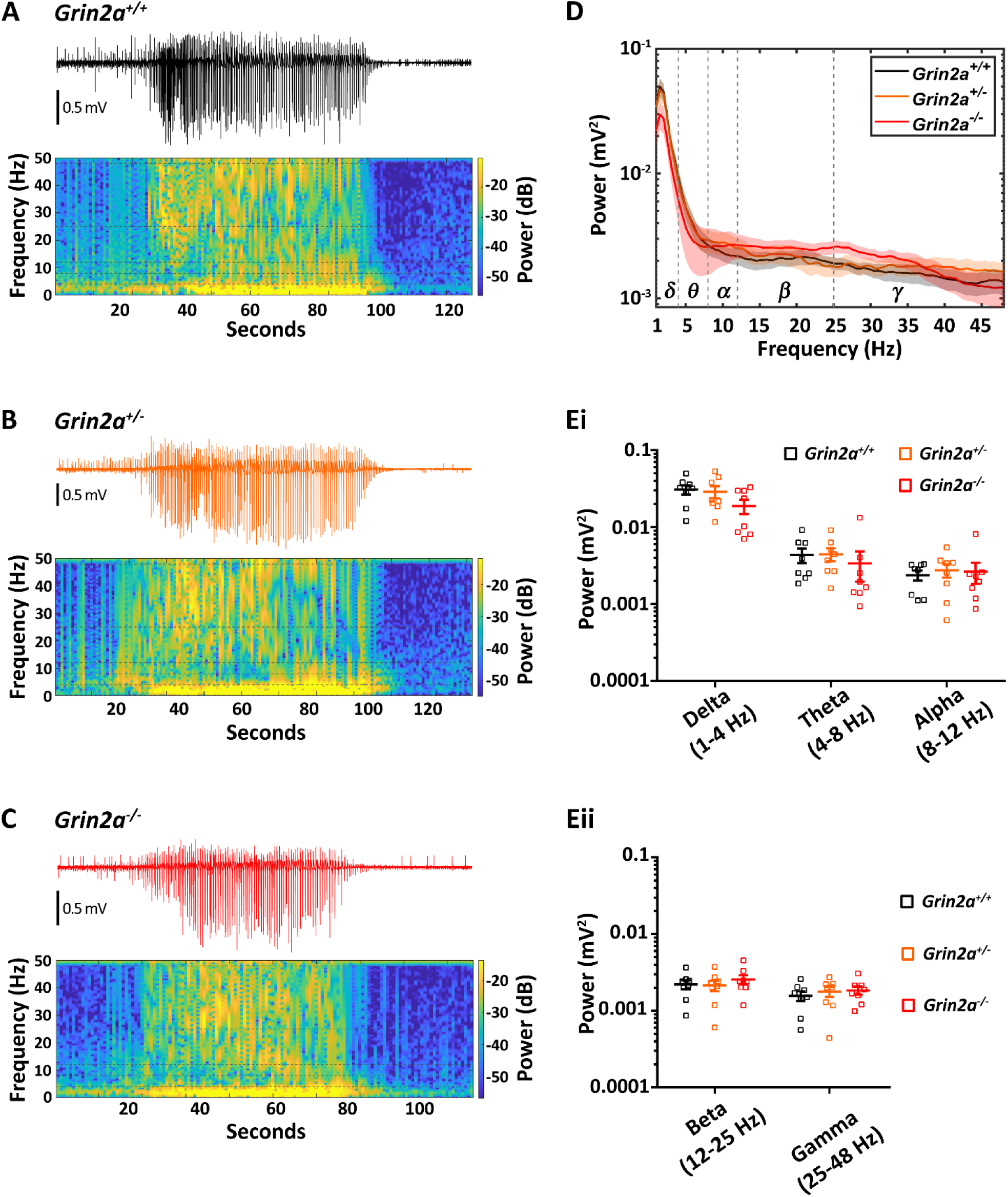
Spectral analysis of SLEs induced by 9.5 mM [K^+^]_e_revealed no genotype-dependent differences between *Grin2a^+/+^, Grin2a^+/-^,* and *Grin2a^-/-^*. (A-C) Representative local field potential (LFP) recordings and associated spectrograms of SLEs from (A) *Grin2a^+/+^,* (B) *Grin2a^+/-^* and (C) *Grin2a^-/-^*. **(D)** Mean power spectral traces for each genotype (solid line: mean; shading: sem). **(Ei, Eii)** Quantification of power in different frequency bands. Delta: Kruskal-Wallis statistic = 3.82, p = 0.15; Theta: Kruskal-Wallis statistic = 3.92, p = 0.14; Alpha: Kruskal-Wallis statistic = 0.79, p = 0.68; Beta: One-way ANOVA, F_2,21_ = 0.45, p = 0.64; Gamma: One-way ANOVA, F_2,21_ = 0.35, p = 0.71. For mean values and multiple comparisons, see Supplementary Table 3B.

## Discussion

In this study, we investigated how reduced GluN2A-NMDA receptor subunit expression influences seizure susceptibility in acute hippocampal slices. Using three mechanistically distinct *in vitro* models of epilepsy, we found that both heterozygous (*Grin2a^+/–^*) and homozygous (*Grin2a^−/−^*) loss of GluN2A reduced the likelihood of seizure-like events (SLEs), particularly in the 0Mg²⁺/5K⁺ and 4-aminopyridine (4-AP) models. One plausible explanation for the reduced seizure susceptibility observed in *Grin2a* null variants is the disruption of NMDA receptor-mediated synaptic currents, integration and thereby reduced ability for dendritic plateau potential generation in excitatory neurons.^21,31,32^ The loss of GluN2A-containing NMDA receptors, coupled with reduced dendritic arborisation, likely diminishes overall excitatory drive and raises the threshold for initiating dendritic plateau potentials, regenerative events that amplify and synchronise excitatory inputs across neuronal networks. This attenuation of dendritic excitability could impair the network dynamics required for ictogenesis.^33,34^ Consistent with this interpretation, recent studies have demonstrated that *Grin2a* deletion leads to altered dendritic morphology and a reduced frequency of miniature excitatory postsynaptic currents in CA1 pyramidal neurons, supporting the idea that GluN2A loss weakens dendritic mechanisms critical for seizure generation.^21^ Furthermore, alterations in inhibitory circuit dynamics could also contribute to reduced seizure susceptibility observed in *Grin2a^+/–^* and *Grin2a^−/−^* networks. Recent findings have demonstrated that *Grin2a* loss leads to impairments in parvalbumin- and somatostatin-positive interneurons, resulting in increased inhibitory input to pyramidal neurons and a shift in the excitatory/inhibitory (E/I) balance towards inhibition^30^. Notably, *Grin2a* loss was associated with increased asynchronous GABA release and elevated gamma-band synchrony, features that could collectively raise the threshold for seizure-like event initiation. Taken together, the combined effects of weakened overall excitatory drive and enhanced or dysregulated inhibition likely act synergistically to suppress ictogenesis in *Grin2a*-deficient networks.

### Distinct Mechanisms of Seizure Generation Reveal Context-Specific Effects of GluN2A Loss

*In vitro* models of epilepsy are invaluable tools for elucidating the mechanisms underlying pathological network hyperexcitability and for screening potential anti-epileptic therapies.^23,35–37^ Different models induce epileptiform activity through distinct mechanisms, and we leveraged this diversity to investigate the specific contributions of GluN2A-containing NMDA receptors to seizure susceptibility and ictogenesis.^23^ In the 4-AP model, 4-aminopyridine preferentially targets voltage-gated K_v_3.1 potassium channels which are predominantly expressed in interneurons, leading to IIEs dominated by GABAergic currents and disinhibition-driven SLEs.^23,38–40^ Its pro-epileptic effects depend heavily on interneuronal firing dynamics.^26,39,41^ GluN2A loss may impair NMDA receptor-mediated excitatory drive onto parvalbumin- and somatostatin-positive interneurons, delaying their recruitment and reducing the emergence of synchronised network discharges, thus resulting in fewer IIEs and delayed SLE onset.^19,20^ In contrast, the 0Mg²⁺/5K⁺ model, which removes the Mg²⁺ block on NMDA receptors and increases glutamatergic drive, produced distinct genotype-dependent differences. *Grin2a^+/–^* and *Grin2a^−/−^* slices exhibited significantly delayed SLE onset and reduced event frequency. This model depends on NMDA receptor-mediated recurrent excitation and dendritic integration – both of which are likely compromised by GluN2A loss, as well as intact feedback and feedforward inhibition.^23,26,28,42^ However, *Grin2a* deficiency has been associated with enhanced inhibitory tone^30^. Together, these changes shift the excitatory/inhibitory balance towards inhibition, which may underlie the reduced susceptibility to seizure-like events in 0Mg^2+^/5K^+^ model. In the high-potassium (high-K⁺) model, seizure generation relies on widespread depolarisation due to elevated extracellular K⁺, which alters intrinsic membrane properties and disrupts inhibitory restraint.^24,43–45^ At 7.5 mM K⁺, *Grin2a^−/−^* slices were less likely to develop SLEs, suggesting a subtle shift in the E/I balance favouring excitation. However, at 9.5 mM K⁺, all genotypes developed robust SLEs, likely due to overwhelming depolarisation that bypasses receptor-specific contributions. This suggests that in strongly depolarising conditions, loss of GluN2A cannot override the non-specific excitatory drive imposed by extracellular K⁺ accumulation. Together, these findings highlight how GluN2A loss differentially influences network excitability and its context-dependent role in modulating seizure susceptibility across mechanistically distinct *in vitro* epilepsy models.

### Implications for *GRIN2A*-related epilepsies

Although many *GRIN2A* mutations identified in patients result in loss-of-function, clinical phenotypes frequently involve hyperexcitability and epilepsy.^9–11,46^ Our findings offer a mechanistic framework to begin reconciling this paradox. GluN2A loss appears to attenuate excitatory drive and reduce susceptibility to SLEs *in vitro*. This finding is consistent with a previous study showing GluN2A antagonism delays the development of seizures *in vivo*.^47^ However, over developmental timescales, changes such as delayed electrographic maturation of fast-spiking parvalbumin-positive interneuron populations, altered synaptic activity, or maladaptive circuit reorganisation may create a substrate for hyperexcitability.^19–22^ Notably, one hypothesis is a compensatory upregulation of GluN2B-containing NMDA receptors, which could prolong excitatory postsynaptic currents, thereby increasing excitability. However, previous studies, including our own, have demonstrated no compensatory increase in GluN2B expression in either *Grin2a^+/-^* or *Grin2a^-/-^* models.^12,21,48^ The absence of such compensation strengthens the argument that reduced GluN2A expression directly diminishes excitatory drive without being offset by increased NMDA receptor function elsewhere. This duality underscores the importance of temporal context when interpreting the consequences of *GRIN2A* mutations: effects that initially reduce network excitability in acute settings may obscure pro-epileptogenic processes that emerge over longer timescales.

While this study provides novel insights into GluN2A-dependent modulation of network excitability, it is limited to acute *in vitro* preparations from young adult rats. No spontaneous seizures were observed in these rat lines during routine husbandry, but subtle phenotypes such as absence seizures or sleep-related discharges cannot be excluded without chronic *in vivo* recordings. Future studies using chronic *in vivo* recordings, optogenetic circuit interrogation, and region-specific manipulations will be essential to dissect how GluN2A loss shapes network function and seizure susceptibility across development and in different brain regions.

In summary, this study demonstrates that reduced GluN2A expression decreases seizure susceptibility in acute hippocampal networks, with implications for dendritic excitability, oscillatory synchronisation, and inhibitory restraint mechanisms. These findings advance our understanding of *GRIN2A*-related epilepsies and support further investigation into the developmental and circuit-level consequences of GluN2A deficiency using *in vivo* and cell-type-specific approaches.

## Data availability

The data that support the findings of this study are available from the corresponding author upon reasonable request.

## Author contributions

NKC, GEH, PCK and DJAW were responsible for the conception and design of the experiments. NKC conducted the electrophysiological recordings; NKC was responsible for the collection, assembly, and analysis of data. NKC, GEH, PCK and DJAW were responsible for the interpretation of data. NKC and DJAW wrote the manuscript, and all authors had the opportunity to contribute to its editing. All persons designated as authors qualify for authorship, and all those who qualify for authorship are listed. All authors read and approved the final manuscript.

## Supplementary Figures and Tables

**Supplemental Figure 1.**
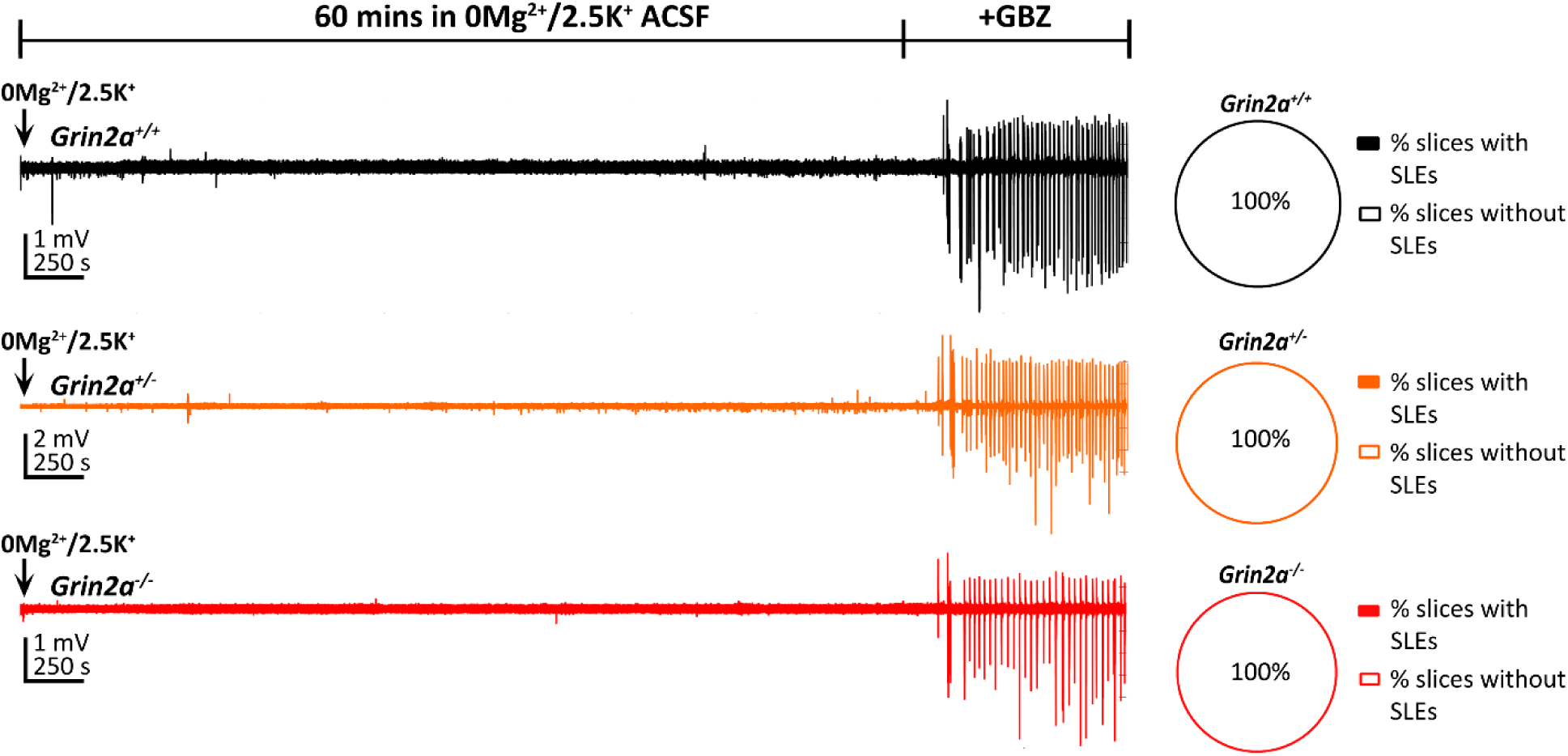
Grin2a^+/+^, Grin2a^+/-^ and Grin2a^-/-^ do not develop SLEs in 0Mg^2+^/2.5K^+^ model. Left: Representative local field potential (LFP) recordings from *Grin2a^+/+^*, *Grin2a^+/-^* and *Grin2a^-/-^* slices bathed in ACSF containing 0 mM magnesium ions and 2.5 mM K^+^ (0Mg^2+^/K^+^) for 1 hour followed by the addition of gabazine (GBZ; 10 μM) to check the viability of slices. Right: Pie graph representing percentage of slices with and without SLEs. Number of slices with SLEs/total number of slices (n rats): *Grin2a^+/+^*, 0/12 (6 rats); *Grin2a^+/-^*, 0/10 (7 rats); *Grin2a^-/-^*, 0/10 (8 rats).

**Supplemental Figure 2.**
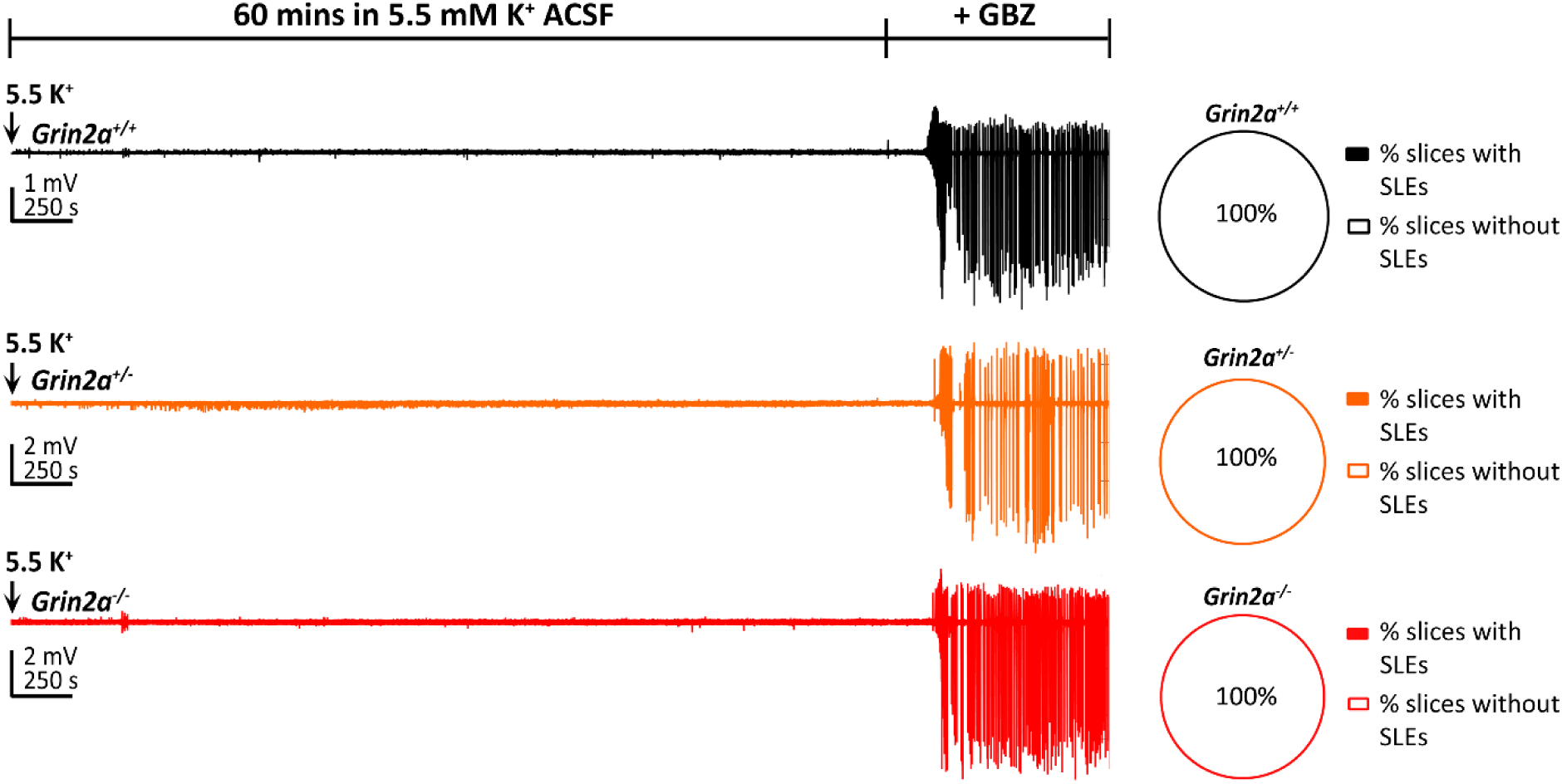
Slices prepared from Grin2a^+/+^, Grin2a^+/-^and Grin2a^-/-^ rats do not develop either IIEs or SLEs in 5.5 mM [K^+^]_e_. Left: Representative local field potential (LFP) recordings from *Grin2a^+/+^, Grin2a^+/-^*and *Grin2a^-/-^* slices bathed in ACSF containing 5.5 mM [K^+^]e for 1 hour followed by the addition of gabazine (GBZ; 10 μM). Right: Pie graph representing percentage of slices with and without SLEs. Number of slices with SLEs / total number of slices (n rats): *Grin2a^+/+^*, 0/10 (6 rats); *Grin2a^+/-^*, 0/10 (6 rats); *Grin2a^-/-^*, 0/10 (6 rats).

**Supplementary Table 1A:**
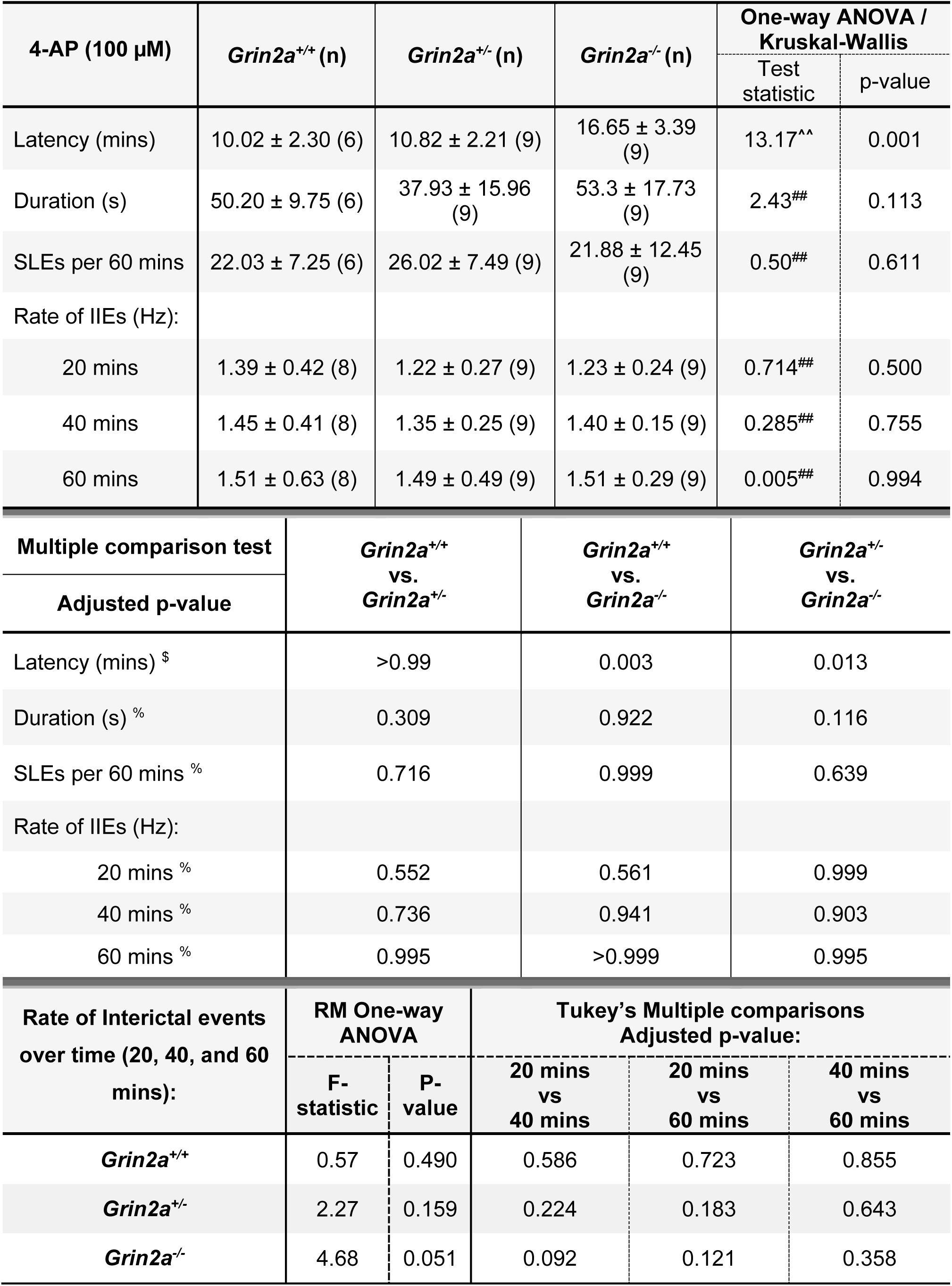
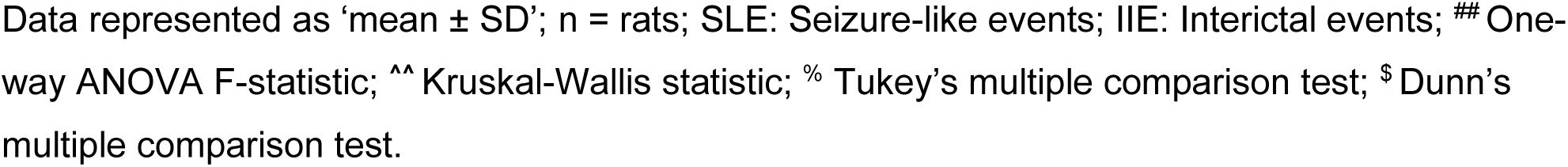
4-Aminopyridine (100 µM) model - SLE and IIE metrics.

| 4-AP (100 μM) | *Grin2a*<sup>+/+</sup> (n) | *Grin2a*<sup>+/-</sup> (n) | *Grin2a*<sup>-/-</sup> (n) | One-way ANOVA / Kruskal-Wallis | |
| --- | --- | --- | --- | --- | --- |
|  |  |  |  | Test statistic | p-value |
| Latency (mins) | 10.02 ± 2.30 (6) | 10.82 ± 2.21 (9) | 16.65 ± 3.39 (9) | 13.17^^ | 0.001 |
| Duration (s) | 50.20 ± 9.75 (6) | 37.93 ± 15.96 (9) | 53.3 ± 17.73 (9) | 2.43## | 0.113 |
| SLEs per 60 mins | 22.03 ± 7.25 (6) | 26.02 ± 7.49 (9) | 21.88 ± 12.45 (9) | 0.50## | 0.611 |
| Rate of IIEs (Hz): |  |  |  |  |  |
| 20 mins | 1.39 ± 0.42 (8) | 1.22 ± 0.27 (9) | 1.23 ± 0.24 (9) | 0.714## | 0.500 |
| 40 mins | 1.45 ± 0.41 (8) | 1.35 ± 0.25 (9) | 1.40 ± 0.15 (9) | 0.285## | 0.755 |
| 60 mins | 1.51 ± 0.63 (8) | 1.49 ± 0.49 (9) | 1.51 ± 0.29 (9) | 0.005## | 0.994 |
| Multiple comparison test | *Grin2a*<sup>+/+</sup> vs. *Grin2a*<sup>+/-</sup> |  | *Grin2a*<sup>+/+</sup> vs. *Grin2a*<sup>-/-</sup> |  | *Grin2a*<sup>+/-</sup> vs. *Grin2a*<sup>-/-</sup> |
| Adjusted p-value |  |  |  |  |  |
| Latency (mins) <sup>\$</sup> | >0.99 | | 0.003 | | 0.013 |
| Duration (s) <sup>%</sup> | 0.309 |  | 0.922 |  | 0.116 |
| SLEs per 60 mins <sup>%</sup> | 0.716 |  | 0.999 |  | 0.639 |
| Rate of IIEs (Hz): |  |  |  |  |  |
| 20 mins <sup>%</sup> | 0.552 |  | 0.561 |  | 0.999 |
| 40 mins <sup>%</sup> | 0.736 |  | 0.941 |  | 0.903 |
| 60 mins <sup>%</sup> | 0.995 |  | >0.999 |  | 0.995 |
| Rate of Interictal events over time (20, 40, and 60 mins): | RM One-way ANOVA |  | Tukey's Multiple comparisons Adjusted p-value: |  |  |
|  | F-statistic | P-value | 20 mins vs 40 mins | 20 mins vs 60 mins | 40 mins vs 60 mins |
| *Grin2a*<sup>+/+</sup> | 0.57 | 0.490 | 0.586 | 0.723 | 0.855 |
| *Grin2a*<sup>+/-</sup> | 2.27 | 0.159 | 0.224 | 0.183 | 0.643 |
| *Grin2a*<sup>-/-</sup> | 4.68 | 0.051 | 0.092 | 0.121 | 0.358 |

- 1 Data represented as 'mean ± SD'; n = rats; SLE: Seizure-like events; IIE: Interictal events; ## One- - 2 way ANOVA F-statistic; ^^ Kruskal-Wallis statistic; % Tukey's multiple comparison test; \$ Dunn's - 3 multiple comparison test.

**Supplemental Table 1B:** 4-Aminopyridine (100 µM) model - SLE power spectral data.

| 4-AP (100 μM) | *Grin2a*<sup>+/+</sup> (n) | *Grin2a*<sup>+/-</sup> (n) | *Grin2a*<sup>-/-</sup> (n) | One-way ANOVA / Kruskal-Wallis | |
| --- | --- | --- | --- | --- | --- |
| Power (x10<sup>-4</sup> mV<sup>2</sup>) |  |  |  | Test statistic | p-value |
| Delta (1-4 Hz) | 239.01 ± 31.87 (6) | 209.87 ± 39.61 (9) | 253.93 ± 47.30 (9) | 0.311<sup>##</sup> | 0.74 |
| Theta (4-8 Hz) | 119.01 ± 21.46 (6) | 77.84 ± 16.68 (9) | 131.40 ± 27.65 (9) | 1.63<sup>##</sup> | 0.22 |
| Alpha (8-12 Hz) | 125.73 ± 10.29 (6) | 99.70 ± 12.85 (9) | 123.37 ± 19.26 (9) | 0.86<sup>##</sup> | 0.44 |
| Beta (12-25 Hz) | 82.15 ± 4.99 (6) | 86.35 ± 8.73 (9) | 73.75 ± 6.51 (9) | 0.83<sup>##</sup> | 0.45 |
| Gamma (25-48 Hz) | 22.01 ± 2.37 (6) | 24.92 ± 3.19 (9) | 19.39 ± 3.98 (9) | 0.71<sup>##</sup> | 0.51 |
| Multiple comparison test | *Grin2a*<sup>+/+</sup> vs. *Grin2a*<sup>+/-</sup> | *Grin2a*<sup>+/+</sup> vs. *Grin2a*<sup>-/-</sup> | *Grin2a*<sup>+/-</sup> vs. *Grin2a*<sup>-/-</sup> |  |  |
| Adjusted p-value |  |  |  |  |  |
| Delta (1-4 Hz) % | 0.885 | 0.973 | 0.722 |  |  |
| Theta (4-8 Hz) % | 0.466 | 0.931 | 0.212 |  |  |
| Alpha (8-12 Hz) % | 0.520 | 0.995 | 0.509 |  |  |
| Beta (12-25 Hz) % | 0.924 | 0.732 | 0.427 |  |  |
| Gamma (25-48 Hz) % | 0.843 | 0.870 | 0.473 |  |  |

2 Data represented as 'mean ± sem'; n = rats; SLE: Seizure-like events; <sup>##</sup> One-way ANOVA F- 3 statistic; <sup>^^</sup> Kruskal-Wallis statistic; % Tukey's multiple comparison test; \$ Dunn's multiple 4 comparison test.

**Supplemental Table 2A:**
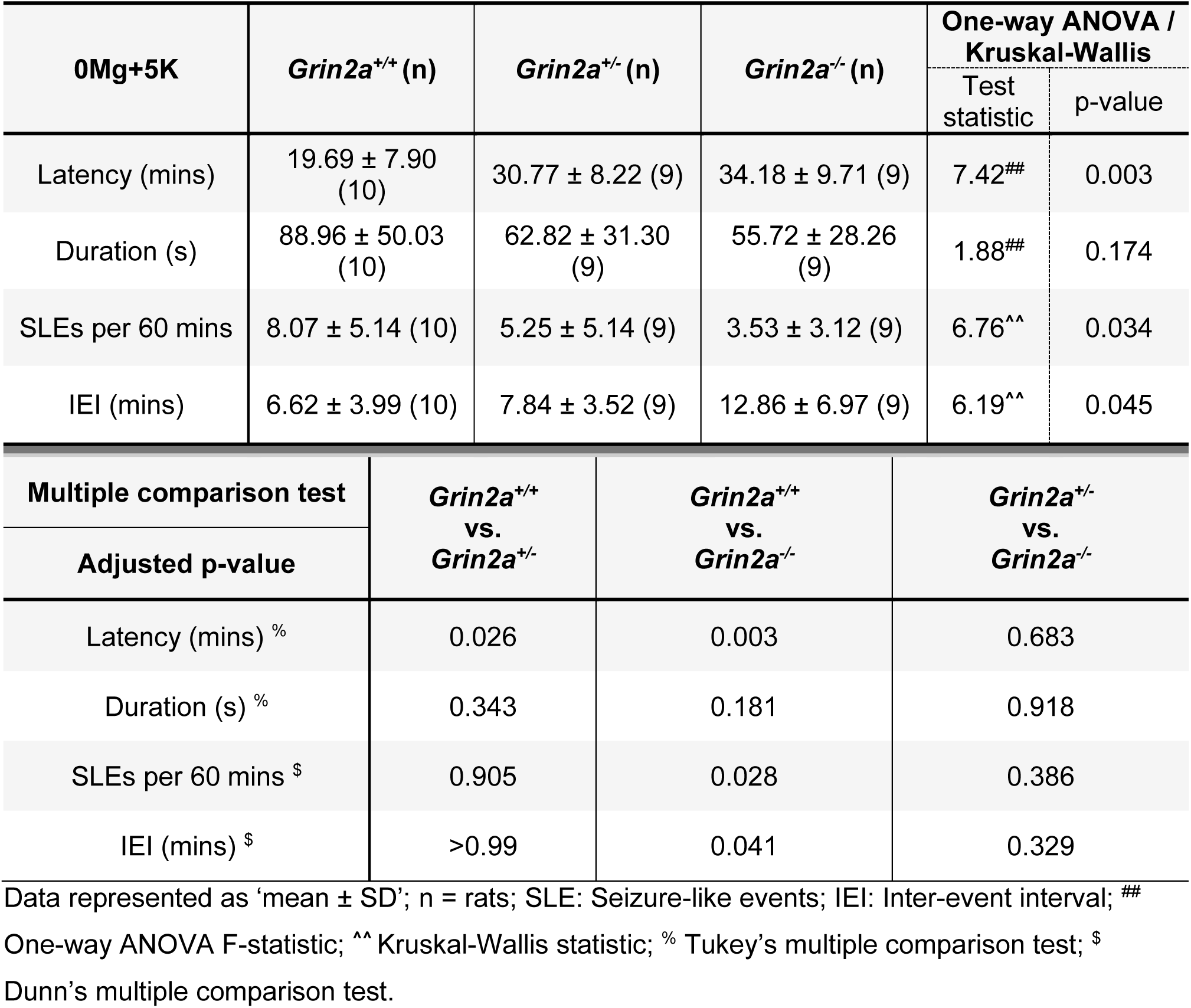
Zero-Magnesium + 5 mM Potassium model – SLE metrics.

| 0Mg+5K | *Grin2a*<sup>+/+</sup> (n) | *Grin2a*<sup>+/-</sup> (n) | *Grin2a*<sup>-/-</sup> (n) | One-way ANOVA / Kruskal-Wallis |  |
| --- | --- | --- | --- | --- | --- |
|  |  |  |  | Test statistic | p-value |
| Latency (mins) | 19.69 ± 7.90 (10) | 30.77 ± 8.22 (9) | 34.18 ± 9.71 (9) | 7.42<sup>##</sup> | 0.003 |
| Duration (s) | 88.96 ± 50.03 (10) | 62.82 ± 31.30 (9) | 55.72 ± 28.26 (9) | 1.88<sup>##</sup> | 0.174 |
| SLEs per 60 mins | 8.07 ± 5.14 (10) | 5.25 ± 5.14 (9) | 3.53 ± 3.12 (9) | 6.76<sup>^^</sup> | 0.034 |
| IEI (mins) | 6.62 ± 3.99 (10) | 7.84 ± 3.52 (9) | 12.86 ± 6.97 (9) | 6.19<sup>^^</sup> | 0.045 |
| Multiple comparison test |  | *Grin2a*<sup>+/+</sup> vs. *Grin2a*<sup>+/-</sup> | *Grin2a*<sup>+/+</sup> vs. *Grin2a*<sup>-/-</sup> | *Grin2a*<sup>+/-</sup> vs. *Grin2a*<sup>-/-</sup> |  |
| Adjusted p-value |  |  |  |  |  |
| Latency (mins) % |  | 0.026 | 0.003 | 0.683 |  |
| Duration (s) % |  | 0.343 | 0.181 | 0.918 |  |
| SLEs per 60 mins \$ | | 0.905 | 0.028 | 0.386 | |
| IEI (mins) \$ | | >0.99 | 0.041 | 0.329 | |

2 Data represented as 'mean ± SD'; n = rats; SLE: Seizure-like events; IEI: Inter-event interval; <sup>##</sup>3 One-way ANOVA F-statistic; <sup>^^</sup> Kruskal-Wallis statistic; % Tukey's multiple comparison test; \$
4 Dunn's multiple comparison test.

**Supplementary Table 2B:** Zero-Magnesium + 5 mM Potassium model – SLE power spectral data.

| 0Mg+5K | *Grin2a*<sup>+/+</sup> (n) | *Grin2a*<sup>+/-</sup> (n) | *Grin2a*<sup>-/-</sup> (n) | One-way ANOVA / Kruskal-Wallis |  |
| --- | --- | --- | --- | --- | --- |
| Power (x10<sup>-4</sup> mV<sup>2</sup>) |  |  |  | Test statistic | p-value |
| Delta (1-4 Hz) | 331.77 ± 34.13 (10) | 321.76 ± 47.84 (9) | 172.91 ± 27.00 (9) | 5.64<sup>##</sup> | 0.009 |
| Theta (4-8 Hz) | 99.75 ± 12.98 (10) | 95.16 ± 19.41 (9) | 35.29 ± 6.09 (9) | 14.46<sup>^^</sup> | <0.001 |
| Alpha (8-12 Hz) | 69.79 ± 8.95 (10) | 59.01 ± 9.97 (9) | 34.85 ± 5.46 (9) | 4.53<sup>##</sup> | 0.02 |
| Beta (12-25 Hz) | 51.73 ± 8.92 (10) | 42.65 ± 7.51 (9) | 61.50 ± 10.54 (9) | 1.03<sup>##</sup> | 0.37 |
| Gamma (25-48 Hz) | 17.43 ± 2.04 (10) | 15.39 ± 1.52 (9) | 31.02 ± 3.01 (9) | 13.75<sup>##</sup> | <0.0001 |

| Multiple comparison test | *Grin2a*<sup>+/+</sup> vs. *Grin2a*<sup>+/-</sup> | *Grin2a*<sup>+/+</sup> vs. *Grin2a*<sup>-/-</sup> | *Grin2a*<sup>+/-</sup> vs. *Grin2a*<sup>-/-</sup> |
| --- | --- | --- | --- |
| Adjusted p-value |  |  |  |
| Delta (1-4 Hz) % | 0.980 | 0.014 | 0.026 |
| Theta (4-8 Hz) \$ | >0.99 | 0.001 | 0.009 |
| Alpha (8-12 Hz) % | 0.637 | 0.017 | 0.134 |
| Beta (12-25 Hz) % | 0.758 | 0.726 | 0.336 |
| Gamma (25-48 Hz) % | 0.799 | <0.001 | <0.001 |

3 Data represented as 'mean ± sem'; n = rats; SLE: Seizure-like events; <sup>##</sup> One-way ANOVA F- 4 statistic; <sup>^^</sup> Kruskal-Wallis statistic; % Tukey's multiple comparison test; \$ Dunn's multiple 5 comparison test.

**Supplemental Table 3A:** High-Potassium (9.5 mM) model - SLE metrics.

| High-K (9.5 mM) | *Grin2a*<sup>+/+</sup> (n) | *Grin2a*<sup>+/-</sup> (n) | *Grin2a*<sup>-/-</sup> (n) | One-way ANOVA / Kruskal-Wallis |  |
| --- | --- | --- | --- | --- | --- |
|  |  |  |  | Test statistic | p-value |
| Latency (mins) | 12.28 ± 4.33 (8) | 13.76 ± 8.01 (8) | 14.25 ± 6.16 (8) | 0.105<sup>^^</sup> | 0.948 |
| Duration (s) | 74.51 ± 25.91 (8) | 84.77 ± 15.82 (8) | 74.91 ± 12.59 (8) | 1.445<sup>^^</sup> | 0.485 |
| SLEs per 60 mins | 18.11 ± 9.53 (8) | 21.34 ± 8.45 (8) | 17.68 ± 7.04 (8) | 0.453<sup>##</sup> | 0.642 |
| IEI (mins) | 4.91 ± 5.68 (8) | 2.53 ± 0.71 (8) | 3.43 ± 2.26 (8) | 0.965<sup>^^</sup> | 0.617 |
| Multiple comparison test |  | *Grin2a*<sup>+/+</sup> vs. *Grin2a*<sup>+/-</sup> | *Grin2a*<sup>+/+</sup> vs. *Grin2a*<sup>-/-</sup> | *Grin2a*<sup>+/-</sup> vs. *Grin2a*<sup>-/-</sup> |  |
| Adjusted p-value |  |  |  |  |  |
| Latency (mins) <sup>\$</sup> | | >0.999 | >0.999 | >0.999 | |
| Duration (s) <sup>\$</sup> | | >0.999 | >0.999 | 0.688 | |
| SLEs per 60 mins <sup>%</sup> |  | 0.727 | 0.994 | 0.664 |  |
| IEI (mins) |  | >0.999 | >0.999 | >0.999 |  |

2 Data represented as 'mean ± SD'; n = rats; SLE: Seizure-like events; IEI: Inter-event interval; <sup>##</sup>3 One-way ANOVA F-statistic; <sup>^^</sup> Kruskal-Wallis statistic; <sup>%</sup> Tukey's multiple comparison test; <sup>\$</sup>
4 Dunn's multiple comparison test.

**Supplemental Table 3B:** High-Potassium (9.5 mM) model - SLE power spectral data.

| High-K (9.5 mM) | *Grin2a*<sup>+/+</sup> (n) | *Grin2a*<sup>+/-</sup> (n) | *Grin2a*<sup>-/-</sup> (n) | One-way ANOVA / Kruskal-Wallis |  |
| --- | --- | --- | --- | --- | --- |
| Power (x10<sup>-4</sup> mV<sup>2</sup>) |  |  |  | Test statistic | p-value |
| Delta (1-4 Hz) | 306.26 ± 41.97 (8) | 287.96 ± 51.14 (8) | 187.52 ± 40.04 (8) | 3.82<sup>^^</sup> | 0.15 |
| Theta (4-8 Hz) | 43.33 ± 9.30 (8) | 44.40 ± 8.46 (8) | 33.86 ± 14.40 (8) | 3.92<sup>^^</sup> | 0.14 |
| Alpha (8-12 Hz) | 23.64 ± 3.52 (8) | 27.45 ± 5.57 (8) | 26.33 ± 8.12 (8) | 0.79<sup>^^</sup> | 0.68 |
| Beta (12-25 Hz) | 21.92 ± 2.95 (8) | 21.43 ± 3.35 (8) | 25.46 ± 3.52 (8) | 0.45<sup>##</sup> | 0.64 |
| Gamma (25-48 Hz) | 15.62 ± 2.32 (8) | 17.69 ± 2.63 (8) | 18.36 ± 2.28 (8) | 0.35<sup>##</sup> | 0.71 |

| Multiple comparison test | *Grin2a*<sup>+/+</sup> vs. *Grin2a*<sup>+/-</sup> | *Grin2a*<sup>+/+</sup> vs. *Grin2a*<sup>-/-</sup> | *Grin2a*<sup>+/-</sup> vs. *Grin2a*<sup>-/-</sup> |
| --- | --- | --- | --- |
| Adjusted p-value |  |  |  |
| Delta (1-4 Hz) <sup>\$</sup> | >0.999 | 0.183 | 0.472 |
| Theta (4-8 Hz) <sup>\$</sup> | >0.999 | 0.359 | 0.198 |
| Alpha (8-12 Hz) <sup>\$</sup> | >0.999 | >0.999 | >0.999 |
| Beta (12-25 Hz) <sup>%</sup> | 0.993 | 0.728 | 0.664 |
| Gamma (25-48 Hz) <sup>%</sup> | 0.819 | 0.706 | 0.978 |

2 Data represented as 'mean ± sem'; n = rats; SLE: Seizure-like events; ## One-way ANOVA F- 3 statistic; ^^ Kruskal-Wallis statistic; % Tukey's multiple comparison test; \$ Dunn's multiple 4 comparison test.

